# Eye movement-related confounds in neural decoding of visual working memory representations

**DOI:** 10.1101/215509

**Authors:** Pim Mostert, Anke Marit Albers, Loek Brinkman, Larisa Todorova, Peter Kok, Floris P. de Lange

## Abstract

The study of visual working memory (VWM) has recently seen revitalization with the emergence of new insights and theories regarding its neural underpinnings. One crucial ingredient responsible for this progress is the rise of neural decoding techniques. These techniques promise to uncover the representational contents of neural signals, as well as the underlying code and the dynamic profile thereof. Here, we aimed to contribute to the field by subjecting human volunteers to a combined VWM/imagery task, while recording and decoding their neural signals as measured by MEG. At first sight, the results seem to provide evidence for a persistent, stable representation of the memorandum throughout the delay period. However, control analyses revealed that these findings can be explained by subtle, VWM-specific eye movements. As a potential remedy, we demonstrate the use of a functional localizer, which was specifically designed to target bottom-up sensory signals and as such avoids eye movements, to train the neural decoders. This analysis revealed a sustained representation for approximately 1 second, but no longer throughout the entire delay period. We conclude by arguing for more awareness of the potentially pervasive and ubiquitous effects of eye movement-related confounds.

**Significance statement:** Visual working memory is an important aspect of higher cognition and has been subject of much investigation within the field of cognitive neuroscience. Over recent years, these studies have increasingly relied on the use of neural decoding techniques. Here, we show that neural decoding may be susceptible to confounds induced by stimulus-specific eye movements. Such eye movements during working memory have been reported before, and may in fact be a common phenomenon. Given the widespread use of neural decoding and the potentially contaminating effects of eye movements, we therefore believe that our results are of significant relevance for the field.

## Introduction

Visual working memory (VWM) is an essential cognitive function necessary for intelligent and flexible behavior. It allows an individual to retain and utilize visual information about the world for a short period of time, even when the original source of that information is no longer available. How this internal retention of information is instantiated by the central nervous system is not fully understood, but has been subject of thorough investigation, and the analytical tools involved have become increasingly sophisticated. A particularly popular analysis tool in order to non-invasively study the neural underpinnings of VWM in humans is multivariate pattern analysis (Haxby et al., 2014; Grootswagers et al., 2016), or neural decoding, in conjunction with neuroimaging techniques. Here we demonstrate that this way of analysis can be vulnerable to confounding effects of item-specific eye movements.

Neural decoding refers to uncovering a latent variable, for instance stimulus identity, from multivariate patterns in neural signals such as those measured by magnetoencephalography (MEG) or functional magnetic resonance imaging (fMRI). This approach has been frequently applied in the study of VWM in order to elucidate where, when and how a memorandum is encoded in the brain. For instance, in a seminal paper, Harrison and Tong (2009) were able to decode the orientation of a memorized grating from visual cortex. More importantly, they were able to do so using a decoder that was trained on separate localizer blocks in which subjects passively perceived gratings. This shows that visual cortex was not only involved in the encoding of the memory item, but did so using a code similar to that of actual perception. Consistently, at about the same time, another study found very similar results (Serences et al., 2009).

Harrison and Tong's (2009) paradigm paved the way for subsequent neural decoding studies that extended it in varying ways. Albers et al. (2013) and Christophel et al. (2015) investigated the representational contents of neural signals while subjects mentally transformed an internal image. Christophel et al. (2017) and Gayet et al. (2017) asked human volunteers to remember geometrical shapes and Foster et al. (2016) investigated memorization of spatial location. The paradigm has also been ported to electrophysiological studies that use MEG or electroencephalography (EEG) (Wolff et al., 2015, 2017; Foster et al., 2016; King et al., 2016). Not only does this allow for investigating the time course of VWM encoding, the high temporal resolution also enables one to characterize the evolution of the underlying neural code (King and Dehaene, 2014; Grootswagers et al., 2016).

The rise of multivariate decoding techniques has opened up new avenues for VWM research. This has led to new insights regarding how VWM is implemented in the brain. First, whereas it has long been known that neurons in the lateral prefrontal cortex (lPFC) are involved in working memory (Curtis and D’Esposito, 2003), it has been unclear how exactly these would encode the working memory item. Neural decoding studies (Harrison and Tong, 2009; Albers et al., 2013) have contributed to the notion that high-fidelity representations may be maintained in the relevant sensory cortex, instantiated by top-down modulation from lPFC neurons (Sreenivasan et al., 2014). Second, a relatively new theory posits that memoranda may not be encoded in an *active* form, i.e. neural spiking, but instead in a *silent* form (Stokes, 2015; Rose et al., 2016; Rademaker and Serences, 2017). According to this hypothesis, memorizing a stimulus establishes a stimulus-specific hidden state throughout the delay interval, possibly via rapid short-term synaptic plasticity. The hidden state then modulates the neural activity generated by a neutral stimulus (e.g. response cue), ultimately triggering a behavioral response contingent on the originally presented item. Crucially, these hidden states do not necessarily evoke activity by themselves and may therefore be challenging to pick up by contemporary neuroimaging techniques (Wolff et al., 2017). Third, there is now increasing evidence that the neural code underlying VWM items is dynamic, changing rapidly in the order of tens of milliseconds, rather than being stable and consistent across time (Stokes et al., 2013; Stokes, 2015; King et al., 2016; Spaak et al., 2017).

It is clear that the study of VWM has seen a proliferation with the advance of decoding methods and that this has resulted in novel, empirically testable theories. The aim of this study was to add to this growing field, by subjecting human volunteers to a combined VWM/imagery task while tracing the representational contents of their neural activity as measured by MEG. While the initial analysis seemed to provide evidence for an active and sustained representation of items in working memory, which was stable throughout a delay interval as long as 8 seconds, follow-up control analyses revealed that this effect could be well explained by stimulus-specific eye movements. These confounding effects may be particularly problematic for studies that employ decoding techniques, owing to their high sensitivity. Given the ubiquity of those techniques, the problem may be pervasive and may require more attention.

## Materials & Methods

### Subjects

Thirty-six human volunteers were recruited from the local institute’s subject pool to participate in a behavioral screening session. Of these, 24 (thirteen male; mean age: 26.8 year, range: 18-60) were selected to participate in the MEG experiment (see *Experimental design and procedure*). Of these 24 selected subjects, three were excluded from MEG analysis due to poor data quality and another four were excluded from the analyses regarding eye movements, because the eye-tracker failed to track the eye reliably in those subjects. The experiment was approved by the local ethics committee (CMO Arnhem-Nijmegen) and conducted according to the guidelines set out by the committee. All participants provided written informed consent and received either monetary compensation or course credits.

### Stimuli

Stimulation was visual and consisted of sinusoidal gratings with a spatial frequency of 1 cycle/°, 80% contrast and one random phase per experimental block. The gratings were masked at an outer radius of 7.5° and an inner aperture radius of 0.7°, and presented on a gray background (luminance: 186 cd/m^2^). Stimuli were generated and presented using MATLAB with the Psychtoolbox extension (Kleiner et al., 2007).

### Experimental design and procedure

The main task was to vividly imagine and remember an oriented grating and, in some conditions, mentally rotate this grating over a certain angle. Each trial began with a dual cue that indicated both the amount (presented above fixation) and the direction (‘>’ for clockwise and ‘<‘ for counterclockwise, presented below fixation) of mental rotation that was to be performed in that trial (Fig. 1). The amount could be either 0°, 60°, 120° or 180°, in either clockwise or counter-clockwise direction, where 0° corresponded to a VWM task. This condition will henceforth be referred to as the VWM condition, and the other three conditions, which corresponded to imagery, as the mental rotation (MR) conditions. This cue lasted for 417 ms, after which a blank screen was shown for another 417 ms. A fixation dot (4 pixels diameter) was present throughout the entire trial, and throughout the entire block. After the blank, a grating was presented for 217 ms that could have either of three orientations: 15°, 75° or 135° (clockwise with respect to vertical). Next, a blank delay period of 8017 ms followed, during which subjects were required to keep the starting grating in mind and, in a subset of trials, mentally rotate it. The delay period was terminated by the presentation of a probe grating for 217 ms, whose orientation was slightly jittered (see *Staircase procedure*) with respect to the orientation that subjects were supposed to have in mind at that moment. Subjects then indicated with a button press whether the probe was oriented clockwise or counterclockwise relative to their internal image. The response period lasted until 2033 ms post probe, after which feedback was given. There were 3 trials per design cell (4 arcs of rotation and 2 directions) per block, resulting in 24 trials per block. In addition, there were two catch trials per block, in which the probe grating was presented at an earlier moment in the delay interval in order to gauge ongoing rotation. All trials were presented in pseudorandomized order. The catch trials were excluded from further analysis, because subjects indicated to find them difficult and confusing.However, subjects indicated to find these trials difficult and confusing, and we decided to exclude them from analysis. In general, each experiment consisted of six experimental blocks (though some subjects performed 5, 7 or 8 blocks), preceded by one or more practice blocks, resulting in a total of 144 experimental trials for most of the participants.

**Figure 1.**
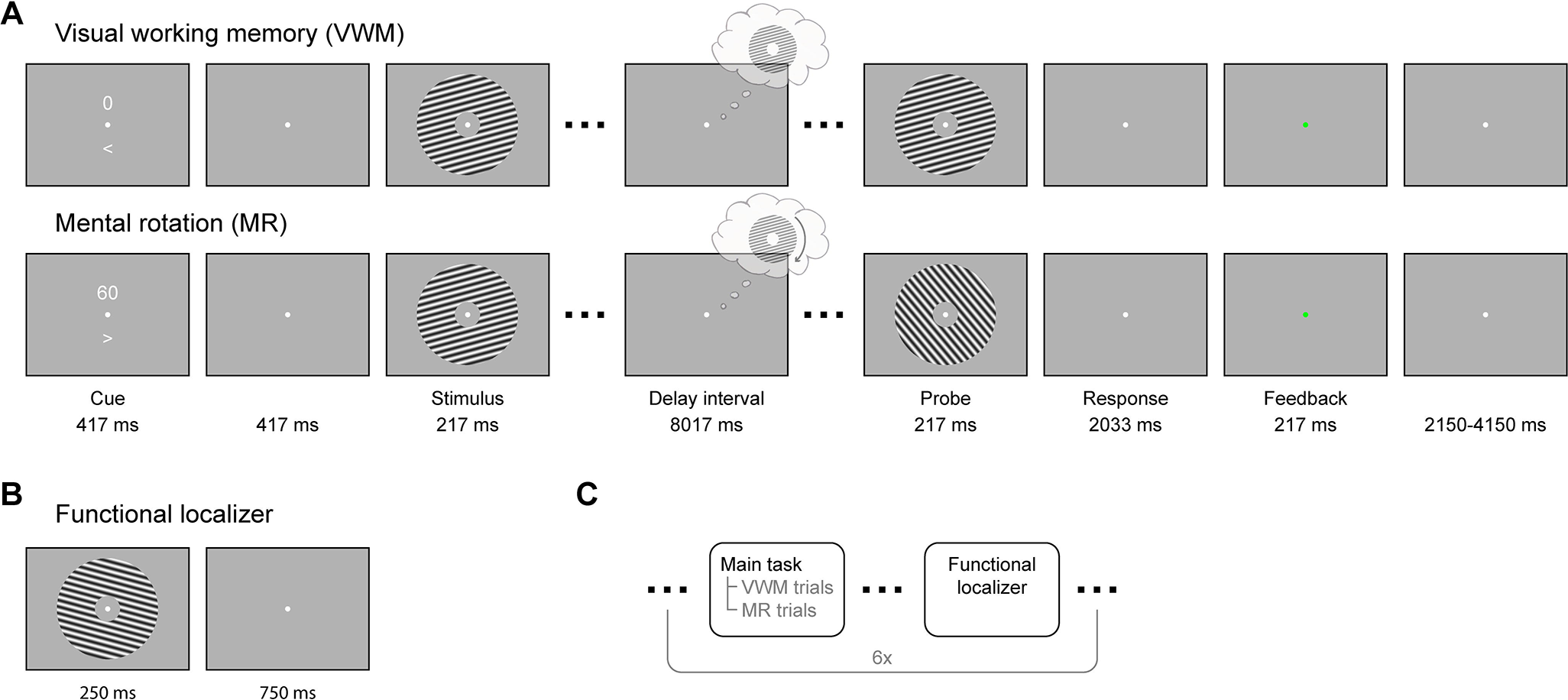
Experimental paradigm. **(A)** In the combined VWM/imagery blocks, subjects were instructed to vividly imagine a grating and to either keep that in mind (VWM condition) or rotate it mentally over a cued number of degrees (MR condition). **(B)** In the functional localizer block, oriented gratings were continuously presented while the subject's attention was drawn to a task at fixation. **(C)** The VWM/imagery and localizer blocks were performed in alternating order.

Interleaved with the VWM/imagery blocks, there were six functional localizer blocks (Fig. 1B, C). In these blocks, gratings of six different orientations (15° to 165°, in steps of 30°) were presented for 250 ms with an inter-trial interval of 750 ms. Each block consisted of 120 trials, resulting in a total of 120 trials per orientation. The task was to press a button when a brief flicker of the fixation dot occurred. Such a flicker occurred between 8 to 12 times (randomly selected number) per block, at random times. Using such a task we ensured that spatial attention was drawn away from the gratings while stimulating subjects to maintain fixation, allowing us to record activity that only reflected bottom-up, sensory-specific signals (Mostert et al., 2015).

Prior to the MEG session, the volunteers participated in a behavioral screening session that served to both train the subjects on the task as well as to assess their capability of carrying it out. Subjects were instructed to mentally rotate the stimulus at an angular velocity of 30 °/s by demonstrating examples of rotations on the screen. Moreover, during this session subjects were required to press a button as soon as they achieved a vivid imagination of the grating upon completion of the cued rotation. This provided a proxy of the speed at which they actually performed the rotation and was used as selection criterion for participation in the MEG session.

### Staircase procedure

The amount of jitter of the probe grating was determined online using an adaptive staircasing procedure to equalize subjective task difficulty across conditions and subjects. The starting difference was set to 15°, and was increased by 1° following an incorrect response and decreased by 0.5488 after two consecutive correct responses. Such a procedure has a theoretical target performance of approximately 80% correct (García-Pérez, 1998). Four separate staircases were utilized, one for each of the VWM and MR conditions.

### MEG recordings, eye-tracker recordings and pre-processing

Neural activity was measured using a whole-head MEG system with 275 axial gradiometers (VSM/CTF Systems, Coquitlam, BC, Canada) situated in a magnetically shielded room. A projector outside the room projected via a mirror system onto the screen located in front of the subject. Fiducial coils positioned on the nasion and in the ears allowed for online monitoring of head position and for correction in between blocks if necessary. Both vertical and horizontal EOG as well as electrocardiogram were obtained to aid in the recognition of artifacts. All signals were sampled at 1200 Hz and analyzed offline using the FieldTrip toolbox (Oostenveld et al., 2010). The data were notch-filtered at 50 Hz and corresponding harmonics to remove line noise, and subsequently inspected in a semi-automatic manner to identify irregular artifacts. After rejection of bad segments, independent component analysis was used to remove components that corresponded to regular artifacts such as heartbeat, blinks and eye movements (although our results suggest that the removal of eye movement-related artifacts was imperfect, see *Results* and *Discussion*). The cleaned data were baseline-corrected on the interval of −200 to 0 ms, relative to stimulus onset.

Gaze position and pupil dilation were continuously tracked throughout the experiment using an Eyelink 1000 (SR Researcher) eye-tracker. The eye-tracker was calibrated before each session and signals were sampled at 1200 Hz. Because we were interested in eye-movements induced by the experimental stimulation, we removed any slow drifts in the signal by baseline-correcting the signal on an interval of −200 to 0 ms relative to cue onset.

### Classification and decoding analyses

Broadly, we conducted two lines of decoding analyses. In the first, we focused only on the blocks in which participants performed the combined VWM/imagery task, using 8-fold cross-validation. We trained a three-class probabilistic classifier that returns the probability that a given trial belongs to either of the three presented grating orientations. In order to improve signal-to-noise ratio, yet retain the ability to draw firm conclusions regarding the timing of any decoded signal, we smoothed the data using a moving average with a window of 100 ms. The classifier was trained across the spatial dimension (i.e. using sensors as features), on trials from all conditions (i.e., all amounts and directions of rotation). This may seem counterproductive, because the mental contents diverge over the delay interval and there should therefore be no systematic relationship between the MEG data and the stimulus label. However, our rationale was that regardless of condition, subjects need to first perceive, encode and maintain the presented stimulus before they can even commence the task, be it VWM or MR. Thus, we expected to be able to extract the neural pattern of the presented stimulus during at least the physical presentation and a brief moment after that. This classifier was then trained and applied across all time points, resulting in a temporal generalization matrix (King and Dehaene, 2014). It is important to note that we trained the classifiers only using the labels of the presented stimulus, but sorted the data in varying ways when testing the performance. For example, by looking at an early training time point, but a late testing time point, we tested whether we could decode the orientation of the grating kept in mind near the end of the delay period, on the basis of the pattern evoked by the presented stimulus early in the trial.

In the second line of analysis, we trained a continuous orientation decoder on the functional localizer and applied this to the VWM/imagery task. We subsequently related the decoded orientation to the true presented orientation by calculating a quantity intuitively similar to a correlation coefficient (see below). Here too, we extended the procedure to include all pairwise training and testing time points, resulting in temporal generalization matrices (King and Dehaene, 2014).

The three-class classifier was based on Bishop (2006, p. 196-199). Briefly, the class-conditional densities were modeled as Gaussian distributions with assumed equal covariance. By means of Bayes’ theorem, and assuming a flat prior, this model was inverted to yield the posterior probabilities, given the data. Specifically, let **x** be a column vector with length equal to the number of features [number of sensors for MEG data, two for gaze position (horizontal and vertical location)] containing the data to be classified, then the posterior probability that the data belongs to class *k* is given by the following equations:

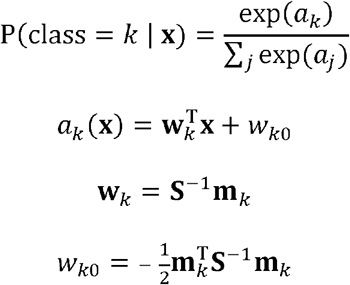

where **m**_*k*_ is the mean of class *k* and **S** is the common covariance, both obtained from the training set. The latter was calculated as the unweighted mean of the three covariance matrices for each individual class, and subsequently regularized using shrinkage (Blankertz et al., 2011) with a regularization parameter of 0.05 for the MEG data and 0.01 for the eye-tracker analysis.

The continuous orientation decoder was based on the forward-modeling approach as described in (Brouwer and Heeger, 2009, 2011) but adapted for improved performance (Kok et al., 2017). The forward model postulates that a grating with a particular orientation activates a number of hypothetical orientation channels, according to a characteristic tuning curve, that subsequently lead to the measured MEG data. We formulated a model with 24 channels spaced equally around the circle, whose tuning curves were governed by a Von Mises curve with a concentration parameter of 5. Note that all circular quantities in the analyses were multiplied by two, because the formulas we used operate on input that is periodic over a range of 360°, but grating orientation only ranges from 0° to 180°. Next, we inverted the forward model to obtain an inverse model. This model reconstructs activity of the orientation channels, given some test data. In this step we departed from Brouwer and Heeger's (2009, 2011) original formulation in two aspects. First, we estimated each channel independently from each other, allowing us to include more channels than there are stimulus classes. Second, we explicitly took into account the correlational structure of the noise, which is a prominent characteristic of MEG data, in order to improve decoding performance (Blankertz et al., 2011; Mostert et al., 2015). For full implementational details, see (Kok et al., 2017). The decoding analysis yields a vector **c** of length equal to number of channels (24 in our case) with the estimated channel activity in a test trial, for each pairwise training and testing time point. These channels activities were then transformed into a single orientation estimate *θ* by calculating the circular mean (Berens, 2009) across all the orientations the channels are tuned for, weighted by each individual activation:

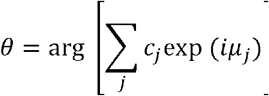

where the summation is over channels, *µ_j_* is the orientation around which the *j*th channel tuning curve is centered, and *i* is the imaginary unit. These decoded orientations can then be related to the true orientation, across trials, as follows:

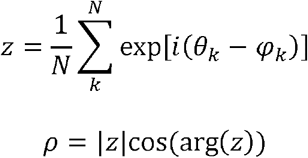

where *N* is the number of trials and *φk* is the true orientation on trial *k*. The quantity *ρ* is also known as the test statistic in the V-test for circular uniformity, where the orientation under the alternative hypothesis is pre-specified (Berens, 2009). This quantity has properties that make it intuitively similar to a correlation coefficient: it is +1 when decoded and true orientations are exactly equal, −1 when they are in perfect counter-phase and 0 when there is no systematic relationship or when they are perfectly orthogonal.

### Statistical testing

All inferential statistics were performed by means of a permutation test with cluster-based multiple comparisons correction (Maris and Oostenveld, 2007). These were applied to either whole temporal generalization matrices, or horizontal cross-sections thereof (i.e. a fixed training window). These matrices/cross-sections were tested against chance-level (33%) in the classification analysis, or against zero in the continuous decoding analysis. In the first step of each permutation, clusters were defined by adjacent points that crossed a threshold of p < 0.05 according to a two-tailed one-sample t-test. The t-values were summed within each cluster, but separately for positive and negative clusters, and the largest of these were included in the permutation distributions. A cluster in the true data was considered significant if its p-value was less than 0.05. For each test, 10,000 permutations were conducted.

### Spatial patterns and source analysis

To interpret the signals that the classifier and decoder pick up, we looked at the corresponding spatial patterns (Haufe et al., 2014). The spatial pattern is the signal that would be measured if the latent variable that is being decoded is varied by one unit. For both the probabilistic classifier and the continuous orientation decoder, this comes down to the difference ERF between each category and the average across all categories. This yields one spatial pattern for each class, and these were subsequently averaged across classes, as well as across time of interest, and fed into planar transformation and source analysis.

For source analysis, we used a template anatomical scan provided by FieldTrip to create a volume conduction model based on a single shell model of the inner surface of the skull. The source model consisted of a regular grid spaced 0.5 cm apart that encompassed the entire brain. Leadfields were calculated and rank-reduced to two dimensions, to accommodate the fact that MEG is blind to tangential sources. The covariance of the data was calculated over the window of 1 to 8 s post-stimulus and regularized using shrinkage (Blankertz et al., 2011) with a regularization parameter of 0.05. The leadfields and data covariance were then used to calculate linearly constrained minimum variance spatial filters (LCMV, also known as beamformers; Van Veen et al., 1997). Applying these filters to sensor-level data yields activity estimates of a two-dimensional dipole at each grid point. We further reduced these estimates to a scalar value by means of the Pythagorean theorem. This leads to a positivity bias however, that we corrected for using a permutation procedure (see Manahova et al., 2017, for details). The number of permutations was 10,000. The final result was interpolated to be projected on a cortical surface, and quantifies the degree to which a particular area contributed to the performance of the classifier/decoder.

## Results

Human volunteers performed a combined VWM/imagery task (Fig. 1A), in which they were presented with an oriented grating (15°, 75° or 135°) that they either kept vividly in mind during the delay interval, or mentally rotated for a certain number of degrees. The amount and direction of rotation were randomized across trials and were indicated at the start of each trial. Subjects were trained in a separate behavioral screening session to perform the mental rotation at a fixed angular velocity, in order to facilitate comparison between conditions as well as across subjects. When the rotation was completed, the subject was instructed to keep the final image in mind for the remainder of the delay interval. After the delay interval, a probe grating was presented, whose orientation was slightly jittered with respect to the orientation the subjects were supposed to have in mind at the end of the interval. The task was to indicate whether the probe was oriented clockwise or counterclockwise with respect to the grating help in mind. The amount of jitter was adapted online using a staircase procedure, separately for each of the four conditions, in order to equalize subjective difficulty.

### Behavioral results

Subjects were selected in a behavioral screening session to ensure that only subjects capable of doing the task would participate in the MEG session. In the MEG session, the average accuracies for the four condition ranged from 68-72%, confirming that subjects were able to do the task, as well as that the staircase procedure was successful. The average final jitter estimate from the staircase procedure for the 0°, 60°, 120° and 180° conditions were as follows (standard error of the mean in parentheses): 3.1° (0.54°), 11.0° (1.23°), 13.4° (1.65°) and 6.5° (1.38°), respectively. With the exception of the 180° condition, the rising trend in these values shows that subjects found the task more difficult when the amount of rotation was larger. The relatively low value for the 180° condition however indicates that this condition was relatively easy. It suggests that subjects either did not perform the half-circle rotation at all, despite being explicitly told to do so, or that they also memorized the starting orientation and used this in their judgment.

### Sustained decoding of VWM items from MEG signals

In order to assess the representational contents of the neural signals while the subjects were engaged in VWM/imagery, we constructed a three-class probabilistic classifier that yields the posterior probabilities that any given data belong to the either of three presented orientations. That is, the classifier was trained according to the labels of the presented stimulus. Furthermore, we applied temporal generalization to investigate the dynamics of the neural pattern (King and Dehaene, 2014). The classifier was trained on trials from all conditions (i.e., all amounts and directions of rotation) pooled together to obtain maximum sensitivity (see *Methods* for rationale). To verify that we could decode stimulus identity, we applied the classifier to the same (pooled) data using cross-validation, and found successful decoding during a period of up to approximately 2.5 seconds after stimulus onset (Fig. 2-1). The stimulus itself was presented for only 250 ms and therefore the later part of this period necessarily indicates an endogenous representation, presumably stemming from active mental instantiation by the subject.

Next we looked at the decoding performance in the VWM condition alone, using the classifier trained on all conditions as described above. We found that the identity of the memory item could be decoded during the entire delay interval, using classifiers obtained from a training window of approximately 0.5-1.5 s (Fig. 2A). The performance stayed above chance-level (33.3%) at a stable level of ~37% throughout the entire interval (Fig. 2B). In contrast, no such sustained generalization was observed for classifiers trained at approximately 0-0.5 s. In other words, the maintenance of the memory item is described by the same neural code as employed in the presumed mental reinstantiation of the stimulus shortly after its presentation, but not by the neural code evoked by the initial presentation of the stimulus.

**Figure 2.**
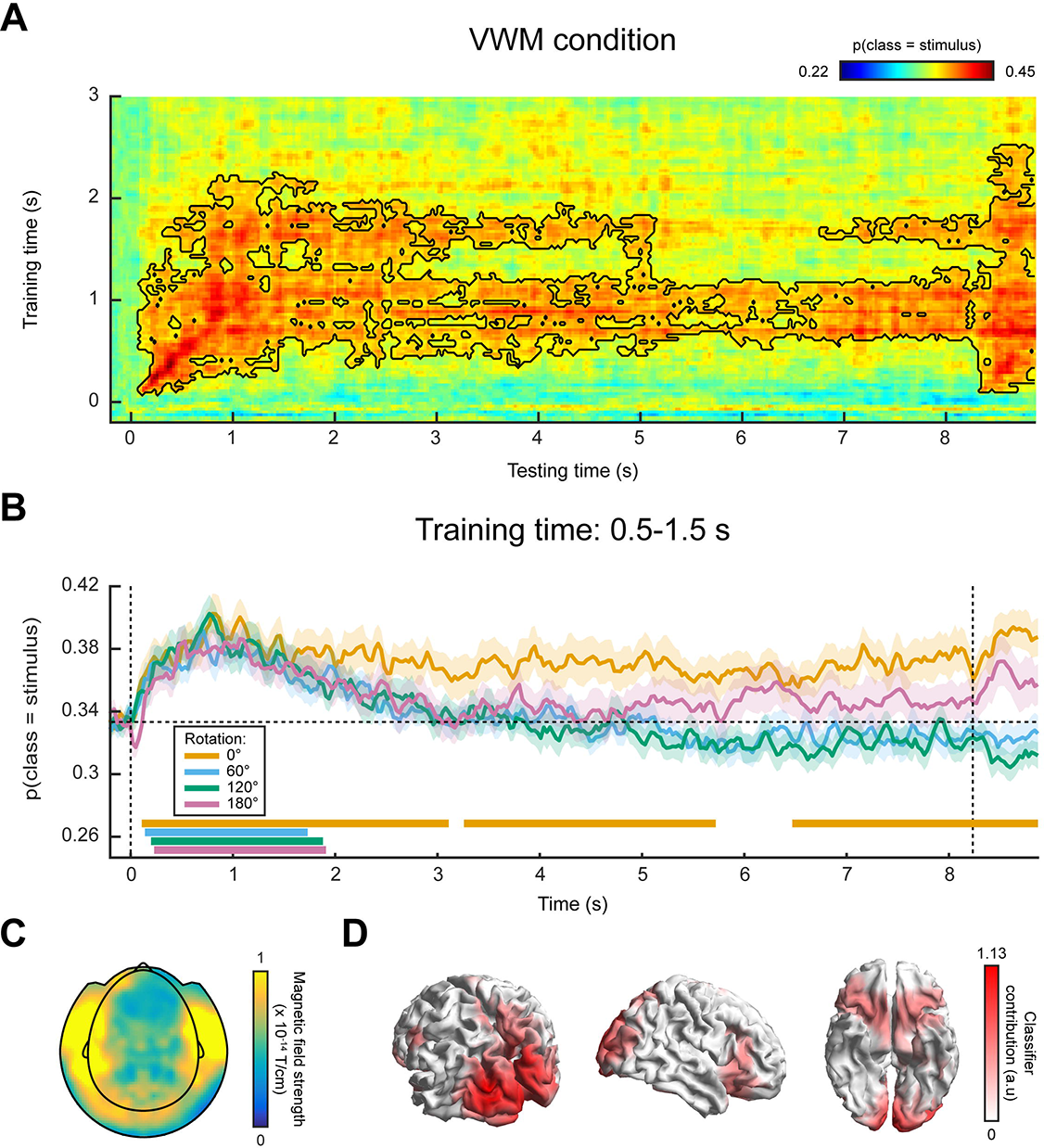
MEG classification results, cross-validation within VWM/imagery task. **(A)** Temporal generalization matrix of classification performance in the VWM condition. The color scale denotes the average posterior probability that the data belong to the same class as the presented stimulus. The black outline demarcates a significant cluster (p = 0.006). **(B)** Classification performance averaged over the training time window of 0.5-1.5 s, separately for the VWM and the three MR conditions. Note that the 0° condition corresponds directly to the matrix in (A). The two vertical dashed lines indicate stimulus and probe onset. Shaded areas indicate the standard error of the mean (SEM). Significant clusters are indicated by the horizontal bars in the lower part of the figure. **(C)** Planar gradiometer and **(D)** source topography of areas that contribute to the classifier. **Figure 2-1: MEG classification performance within the VWM/imagery task, pooled across VWM and MR conditions.** Temporal generalization matrix of the average posterior probability that the data belong to the same class as the presented stimulus. The black outline corresponds to a significant cluster (p = 0.007). **Figure 2-2: Complete MEG classification results.** A nalogously to Fig. 2, the classifier was trained on the time window of 0.5-1.5 s post stimulus onset, and according to the labels of the presented stimulus. **(A)** Average posterior probability that the data belong to the class of the target orientation, that is, the orientation that the subjects were supposed to have in mind at the end of the delay period. For both the 0° and the 180° conditions, the target orientation was the same as the presented stimulus. For the 60° and 120° conditions however, the target and presented stimulus were different, hence the below-chance probabilities at the beginning of the delay period. **(B)** The average posterior probabilities that the data belong to either of three classes: the same orientation as the presented stimulus, the orientation of the presented stimulus ±60° or the orientation of the presented stimulus ±120°, plotted separately for the four VWM/MR conditions. The plus/minus-sign is due to the mental rotation being performed either clockwise or counter-clockwise. This figure gives insight into whether the representations of any intermediate orientations become active during mental rotation, which is particularly relevant for the 120° and 180° conditions. For instance, in the 180° counter-clockwise condition, the subject would start with a mental image with the same orientation as the presented stimulus, then pass through respectively −60° and −120°, ultimately to reach the target of −180° (i.e. 0°). If the neural representations of all these orientations become active in sequence, one would first expect a peak in posterior probability that the data belong to the same class as the stimulus (gray line, bottom figure), then a peak in the probability of belonging to the presented stimulus −60° (blue line), then for −120° (orange line) and finally again for 0° (gray line). Note that **(A)** and **(B)**, as well as Fig. 2 all depict the same data, but visualized in different manners. Shaded areas denote the SEM and significant clusters are depicted by the thick horizontal lines at the bottom of the panels.

Contrary to previously used paradigms, where two stimuli were displayed at the beginning of a trial and a retro-cue signaled the item that was to be remembered (e.g. Harrison and Tong, 2009; Albers et al., 2013; Christophel et al., 2015, Christophel 2017), in the present experiment we only showed one stimulus. It is therefore possible that our sustained decoding performance is the result of a longer lasting stimulus-driven effect (e.g. stimulus aftereffect), rather than a manifestation of VWM. However, if this were true, then we should find a similar effect in the three MR conditions. If, on the other hand, the classifiers picked up the item held in mind, then the probability that a trial is assigned to the same class as the presented stimulus should drop over time, as the subject rotates the mentally imagined grating away from the starting orientation. Our results demonstrate that the latter is indeed the case (Fig. 2B). Whereas the probability that the data belong to the same class as the presented stimulus stays steadily above chance in the VWM condition, it drops to (below) chance-level in the 60° and 120° conditions. In the 180° condition, the probability initially falls to chance as well, but reemerges later as a rising, though non-significant trend. This is to be expected, because the final orientation that the subjects should have in mind in the 180° condition is identical to the orientation of the presented stimulus at the start of a trial.

In order to facilitate interpretation of these results, we inspected the classifier’s corresponding sensor topography (Fig. 2C) and source localization (Fig. 2D), averaged over the training time period of 0.5-1.5 s. These indicate that both occipital and prefrontal sources contribute to the classifier’s performance.

In the MR conditions, it is not only of interest whether the representation of the initially presented stimulus fades (as the subject mentally rotates it), but also whether the intermediate (for the 120° and 180° conditions) and final orientations can be decoded from the neural signals. We found some indication that the final orientation – but not the intermediate ones (Fig. 2-2B) – indeed emerges halfway through the delay period, but this effect was not statistically significant (Fig. 2-2A).

### Gaze position tracks VWM contents

As a control analysis, we examined the position of subject’s gaze, and in particular whether there was a systematic relationship with the identity of the mental image. It is a well-documented phenomenon that eye movements occur spontaneously during visual imagery in a manner systematically related to the mental image (Brandt and Stark, 1997; Spivey and Geng, 2001; Laeng and Teodorescu, 2002; Laeng et al., 2014; Bone et al., 2017). Although in our experiment the volunteers were instructed to fixate, systematic eye movements in working memory tasks have been observed before (Foster et al., 2016) and we therefore also investigated this possibility given that even minute, involuntary eye-moments could have serious consequences for the interpretation of our results described above (see *Discussion*).

We performed the same analysis as described in the previous section, but now used the gaze position (horizontal and vertical dimensions) as measured by the eye-tracker, instead of the MEG data, as features for the classifier. First, when looking at all trials collapsed across conditions, we found above-chance decoding for stimulus identity in a time period of approximately 0.5-3.5 s post-stimulus that was marginally significant (Fig. 3-1). Then, when looking at the decoded signal within the VWM condition only, we found a sustained pattern, similar to the MEG decoding results (Fig. 3A), though again only marginally significant. This suggests that upon perceiving and encoding the stimulus, subjects move their eyes in a way systematically related to the identity of the stimulus, and keep that gaze position stable throughout the entire delay period.

**Figure 3.**
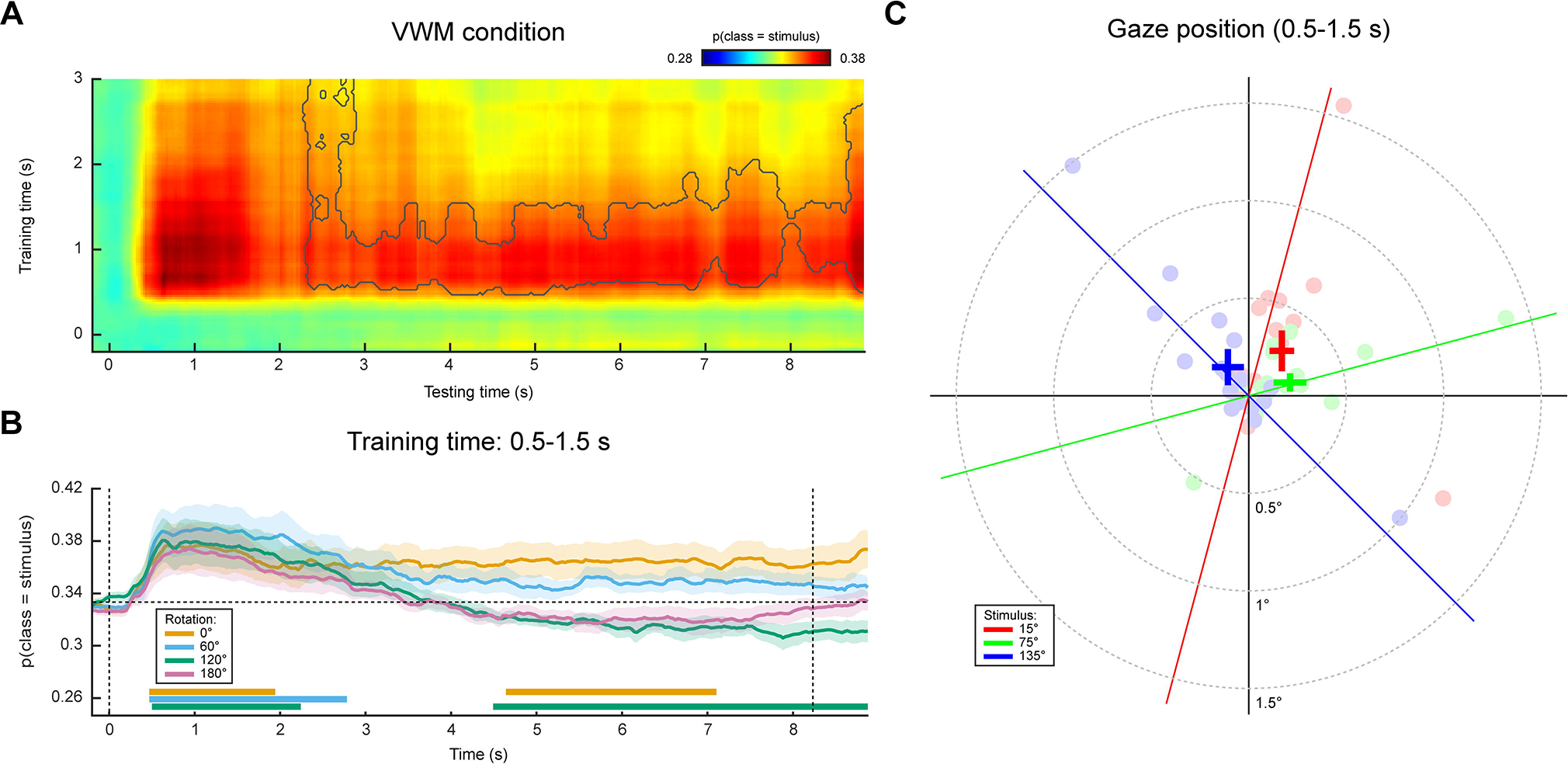
Gaze position classification results. **(A)** Same as in Fig. 2A, except the classification was performed on gaze position as measured by an eye-tracker, rather than on MEG data. The gray outline demarcates a near-significant cluster (p = 0.069). **(B)** Same as in Fig. 2B, except the analysis was performed on gaze position. **(C)** Average gaze position during 0.5-1.5 s after stimulus onset, separately per stimulus orientation. Each transparent dot corresponds to an individual subject. The crosses are the grand averages, where the vertical and horizontal arms denote the SEM. The three colored lines depict the orientation of the three stimuli. **Figure 3-1: Gaze position classification performance within the VWM/imagery task, pooled across VWM and MR conditions.** Similar to Supplementary Fig. 1, except the classifier is trained and tested on gaze position rather than MEG data. Gray outline indicates a near-significant cluster (p = 0.068). **Figure 3-2: Complete gaze position classification results, visualized in a variety of ways.** This figure is analogous to Supplementary Fig. 2, except the classifier is trained and tested on gaze position rather than on MEG data.

Again, we found this sustained pattern to be specific to the VWM condition, because the probability that the data belongs to the same class as the presented stimulus drops over time in the three MR conditions (Fig. 3B). As explained above, this indicates that the sustained above-chance classification in the VWM condition cannot be explained solely by the stimulus itself, but must also reflect the mental image to at least some degree. In fact, in this analysis we even found evidence that the gaze moves towards a position consistent with the orientation of the presented grating plus or minus 60° (depending on the cued direction of rotation) in the MR conditions, but not any further (Fig. 3-2A,B). In short, there was a systematic relationship between gaze position and stimulus orientation, after which the gaze position tracked the orientation kept in mind during the delay period, but only for a maximum of approximately ±60° relative to starting orientation.

Fig. 3C displays the grand average, as well as individual average gaze positions during 0.5-1.5 s after stimulus onset, separately for each of the three stimulus conditions, collapsed across VWM and MR conditions. Although there is large variability among subjects in the magnitude of the eye movements, a general trend can be discerned where subjects position their gaze along the orientation axis of the grating. The mean disparity in visual angle with respect to pre-trial fixation was 0.23°, which is in the same order of magnitude as reported previously (Foster et al., 2016), though for some subjects it was as large as 1.5°.

These findings raise the concern that the sustained decoding of VWM items from MEG signals are the result of stimulus-related eye confounds (see *Discussion* for possible underlying mechanisms). In order to counter this possibility and assess whether the mental image is genuinely encoded in the neural signals, we made use of a functional localizer in a between-task generalization procedure.

### Decoding sensory-specific signals using a functional localizer

Besides the combined VWM/imagery task, subjects also performed a functional localizer task in interleaved blocks (Fig. 1B,C). In this task, subjects were continuously presented with gratings while they performed a challenging detection task at fixation in order to draw the subject’s attention away from the gratings. This task ensured that activity patterns mainly reflected automatic sensory processing of the stimuli and, in addition, discouraged eye movements in response to the stimuli. We used these data for between-task generalization (King and Dehaene, 2014), whereby we trained a continuous orientation decoder on the functional localizer and subsequently applied it to the data from the VWM/imagery task. The chief advantage of this method is that it ensures that the decoder is sensitive to sensory signals only, and not to higher-level top-down processes involved in mentally manipulating an image. It thus allows us to track sensory-specific activation throughout the delay period (Mostert et al., 2015). Cross-validation within the functional localizer confirmed that we were indeed able to reliably decode orientation-specific information from activity evoked by passively perceived gratings (Fig. 4-1). Moreover, we were not able to decode grating orientation on the basis of gaze position, verifying that the data from the functional localizer were not contaminated by stimulus-specific eye movements (Fig. 4-2).

**Figure 4.**
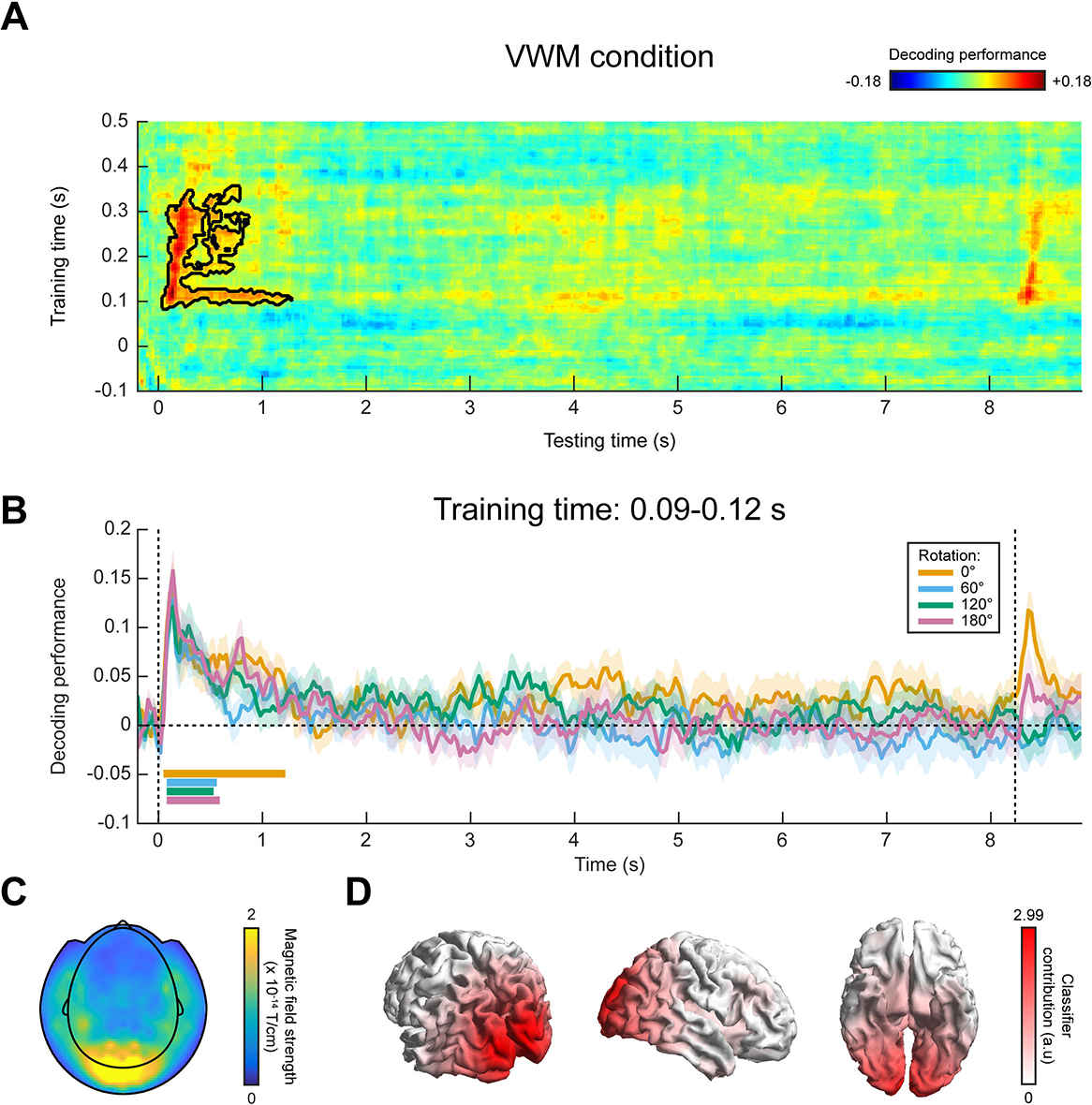
MEG decoding results, generalized from localizer to VWM/imagery task. **(A)** Temporal generalization matrix of orientation decoding performance, for which the decoder was trained on all time points in the functional localizer and tested across all time points in the VWM/imagery task. The color scale reflects the correspondence between true and decoded orientation. The black outline shows a significant cluster (p = 0.04). Note that the x- and y-axis in the figure are differently scaled for optimal visualization. **(B)** Decoding performance over time in the VWM/imagery task, averaged over decoders trained in the window of 0.09-0.12 s in the localizer, separately for the VWM and three MR conditions. Shaded areas denote the SEM and significant clusters are indicated by the horizontal bars. **(C)** Planar gradiometer and **(D)** source topography of areas that contribute to the decoder. **Figure 4-1: MEG decoding performance within the functional localizer.** The time axis represents matched training and testing time-points. Shaded areas denote the SEM, and the horizontal line demarcates a significant cluster (p = ±0). **Figure 4-2: Gaze position classification performance within the functional localizer, using cross-validation. (A)** The time axis represents matched training and testing time-points. No significant above-chance classification was found. Note that although there appears to be a rise in performance after approximately 500 ms, this only reached a value of 0.1675 at its peak (t = 0.51 s), which is very little above chance (0.1667 for six classes). Shaded areas denote SEM. **(B)** Average gaze position at 0.51 s after stimulus onset, separately per stimulus orientation. Each transparent dot corresponds to an individual participant. The crosses are the grand averages, where the vertical and horizontal arms denote the SEM. The six colored lines depict the orientation of the six stimuli.

In the VWM condition, the decoders trained on the functional localizer data could reliably decode the orientation of the presented stimulus (Fig. 4A). Moreover, for a training time of approximately 90-120 ms, we could decode the stimulus for a prolonged time, lasting over 1 s after stimulus onset. Interestingly, this training time point coincides with the time at which peak performance is obtained within the localizer itself (Fig. 4-1). Comparing the decoding trace within this training time window with the three MR conditions, it can be seen that grating orientation can be decoded in all four conditions for a sustained period of approximately 500 ms (Fig. 4B). Furthermore, it is noteworthy that decoding performance in the VWM condition remains above baseline throughout large portions of the interval, although this was not significant.

We inspected the spatial pattern (Fig. 4C) and corresponding source topography (Fig. 4D) for the decoders, averaged across training time 90-120 ms. These highlight primarily occipital regions as contributing to the decoder’s performance, consistent with our premise that the functional localizer primarily induced bottom-up sensory signals, especially during this early time interval (Mostert et al., 2015).

In summary, our findings suggest that the stable, persistent representation found in our within-task MEG decoding result may well be attributed to stimulus-specific eye movements. In contrast, no clear evidence was found for such a long-lasting representation when training the decoder on the functional localizer. Given that the localizer was not contaminated by stimulus-specific eye movements, these results thus provide a more reliable picture of the sensory representations during the delay interval.

## Discussion

Visual working memory, and especially its implementation in the brain, is an intensely studied topic that has recently undergone innovations in the form of new theories (Sreenivasan et al., 2014; Stokes, 2015; Rose et al., 2016; Rademaker and Serences, 2017). These theories revolve around three central issues: whether the identity of the memory items is encoded in parietal and prefrontal cortex or in sensory cortex; whether the items are encoded by means of persistent activity or in hidden, activity-silent states and whether the neural code is dynamic or static. At first sight, our results seem to provide evidence for a stable signal, possibly involving both occipital and prefrontal cortex, which showed persistent encoding of stimulus information throughout the delay period. However, control analysis revealed that very similar results could be obtained by considering gaze position only. This indicates that our MEG results are likely confounded by stimulus-specific eye movements, which jeopardizes the ability to draw firm conclusions about the neural underpinnings of VWM.

There are a least three possible mechanisms via which stimulus-specific eye movements may confound our results. First, eye movements are known to cause stereotypical artifacts in MEG recordings. Due to the positively charged cornea and negatively charged retina, the eyeball acts as a dipole that is picked up by the MEG sensors. The spatial pattern that the dipole evokes is directly related to its rotation, or in other words, to the position of the subject’s gaze (Plöchl et al., 2012). Thus, if the subject moves their eyes in response to the grating in a manner related to the orientation of that grating, then this will induce a specific pattern in the MEG signals, which in turn is directly related to the grating orientation. A decoding analysis applied to these signals is then likely to pick up the patterns evoked by the eyeball dipoles, confounding potential orientation-related information stemming from genuine neural sources. In fact, our source analysis hints at this scenario (Fig. 2D), as the contributions from presumed prefrontal sources closely resemble an ocular source.

Second, if the eyes move, then the projection falling on the retina will also change, even when external visual stimulation remains identical. Thus, if gaze position is systematically modulated by the image that is perceived or kept in mind, then so is the visual information transmitted to the visual cortex. For example, if a vertical grating is presented and kept in VWM, then the subject may subtly move her or his gaze upward. Correspondingly, the fixation dot is now slightly below fixation, thus leading to visual cortex activity that is directly related to the retinotopic position of the fixation dot. Our decoding analysis may thus actually decode the position of the fixation dot, rather than grating orientation, potentially leading to an incorrect conclusion. Source analysis would in this scenario also point to occipital sources, similarly to what we found (Fig. 2D). Note that this mechanism is not specifically dependent on the presence of a fixation dot. A systematic difference in eye position will also lead to changes in the retinotopic position of, for instance, the presentation display or the optically visible part of the MEG helmet.

Third, if gaze position covaries with the mental image, then decoding of the mental image will also reveal areas that encode eye gaze position, such as oculomotor regions in parietal and prefrontal cortex.

Our findings raise the question of why there were task-induced eye movements that were directly related to the grating kept in VWM. In fact, there is a considerable mass of literature that describes the role of eye movements in mental imagery. It has been found that subjects tend to make similar eye movements during imagery as during perception of the same stimulus (Brandt and Stark, 1997; Laeng and Teodorescu, 2002; Laeng et al., 2014). Already proposed by Donald Hebb (Hebb, 1968), it is now thought that eye movements serve to guide the mental reconstruction of an imagined stimulus, possibly by dwelling on salient parts of the image (Spivey and Geng, 2001; Laeng et al., 2014). Moreover, the specificity of the eye movements is also related to neural reactivation (Bone et al., 2017) and recall accuracy (Laeng and Teodorescu, 2002; Laeng et al., 2014; Bone et al., 2017). Our findings are in accordance with these studies. Subjects’ gaze was positioned along the orientation axis of the grating - that is, the visual location within the stimulus that provided the highest information regarding its orientation, and is thus exactly what one would expect given that the task was to make a fine-grained orientation comparison with a probe grating. Importantly however, subjects were explicitly instructed to maintain fixation throughout the entire trial. We nevertheless observed that not all subjects adhered to this requirement, albeit involuntarily.

Despite these problems associated with the systematic eye movements in our experiment, it is still possible that our decoding results do in reality stem from genuine orientation information encoded in true neural sources. In fact, we used independent component analysis in our pre-processing pipeline to (presumably) remove eye-movement artifacts. However, it would be very difficult, if not impossible, to convincingly establish that no artifacts remain and, considering the similarities between the decoding results from the MEG data (Fig. 2A,B) and the gaze position (Fig. 3A,B), we feel any attempts at this would be unwarranted.

Given the potential pervasiveness of systematic eye movements in VWM/imagery tasks, and the demonstrated susceptibility of our analysis methods to these confounds, one wonders whether other studies may have been similarly affected. Clearly, the first mechanism described above involving the eyeball dipole would only affect electrophysiological measurements like electroencephalography and MEG, and has indeed been a concern in practice (Foster et al., 2016). The second mechanism however, whereby stimulus identity is confounded with the retinal position of visual input, would also affect other neuroimaging techniques such as fMRI. This confound could be particularly difficult to recognize, because it would also affect occipital sources. Moreover, because eye movements during imagery have been found to be positively related to performance (Laeng and Teodorescu, 2002), this could potentially explain correlations between VWM decoding and behavioral performance (Wolff et al., 2017). The third mechanism, whereby one directly decodes gaze position from motor areas, could be a problem especially for fMRI which, thanks to its high spatial resolution, might be well able to decode such subtle neural signals. This concern may be especially relevant for studies that investigate the role of areas involved in eye movements or planning thereof, such as frontal eye fields or superior precentral sulcus, in the maintenance of working memory items (Jerde et al., 2012; Ester et al., 2015; Christophel et al., 2017)

This leaves the question of how to deal with eye movements in VWM/imagery tasks. Naturally, it is important to record eye movements during the experiment, for instance using an eye-tracker or electrooculogram (EOG). One can then test for any systematic relationship and, if found, investigate whether it could confound the main results. In our case, for example, decoding of gaze position leads to strikingly similar results as those obtained from the MEG data. Foster et al. (2016) on the other hand found that decoding performance of working memory items decreased throughout the trial, whereas the deviation in gaze position increased, suggesting that eye confounds cannot explain the main findings. Anoth er approach might be to design the experimental task in such a way that eye movements are less likely. For example, by presenting gratings laterally (e.g. Pratte and Tong, 2014; Ester et al., 2015; Wolff et al., 2017), and assuming that VWM items are stored in a retinotopically specific manner (Pratte and Tong, 2014), the involuntary tendency to move one's eyes subtly along the remembered grating's orientation axis may become less strong, because those gratings are located distantly from the gaze's initial location (i.e. central fixation). Finally, a powerful approach could be to adopt a separate functional localizer, which allows specific decoding of functionally defined representations such as bottom-up, sensory-specific signals (Harrison and Tong, 2009; Serences et al., 2009; Albers et al., 2013; Mostert et al., 2015). If the localizer is well designed and not systematically contaminated by eye movements, then eye movements in the main task cannot have a systematic effect on the decoded signal, thus effectively filtering them out.

Using such a functional localizer, we indeed obtained MEG decoding results that were very dissimilar from those obtained on the basis of gaze position. We no longer found persistent activation of an orientation-specific representation throughout the entire delay period. Nevertheless, the sensory pattern did remain above baseline for a period of approximately 1 second, which is relatively long considering that the stimulus was presented for only 250 ms. One explanation is that the stimulus was relevant for the task. Previous work has shown that task relevance may keep the sensory representation online for a prolonged period even after the stimulus is no longer on the screen (Mostert et al., 2015).

In summary, we demonstrate a case where decoding analyses in a VWM/imagery task are heavily confounded by systematic eye movements. Given the high potential benefit of decoding analyses and its widespread use in the study of working memory and mental imagery, we argue that this problem may be more pervasive than is commonly appreciated. We conclude that eye movement confounds should be taken seriously in both the design as well as the analysis phase of future studies.

## Acknowledgements

This work was supported by The Netherlands Organisation for Scientific Research and the European Research Council (PM: NWO Research Talent grant 406-13-001; AMA & LB: NWO Brain and Cognition grant: 433-09-248; PK: NWO Rubicon grant 446-15-004; FPL: NWO Vidi grant 452-13-016, ERC starting grant: 678286, NWO Brain and Cognition grant: 433-09-248). The authors are grateful to Ivan Toni for useful suggestions during the design phase.

## References

Albers AM, Kok P, Toni I, Dijkerman HC, de Lange FP (2013) Shared Representations for Working Memory and Mental Imagery in Early Visual Cortex. Curr Biol 23:1427–1431.

Berens P (2009) CircStat: A MATLAB Toolbox for Circular Statistics. J Stat Softw Vol 1 Issue 10 2009 Available at: https://www.jstatsoft.org/v031/i10.

Bishop CM (2006) Pattern recognition and machine learning. springer.

Blankertz B, Lemm S, Treder M, Haufe S, Müller K-R (2011) Single-trial analysis and classification of ERP components — A tutorial. NeuroImage 56:814–825.

Bone MB, St-Laurent M, Dang C, McQuiggan DA, Ryan JD, Buchsbaum BR (2017) Eye-movement reinstatement and neural reactivation during mental imagery. bioRxiv:107953.

Brandt SA, Stark LW (1997) Spontaneous Eye Movements During Visual Imagery Reflect the Content of the Visual Scene. J Cogn Neurosci 9:27–38.

Brouwer GJ, Heeger DJ (2009) Decoding and Reconstructing Color from Responses in Human Visual Cortex. J Neurosci 29:13992–14003.

Brouwer GJ, Heeger DJ (2011) Cross-orientation suppression in human visual cortex. J Neurophysiol 106:2108–2119.

Christophel TB, Allefeld C, Endisch C, Haynes J-D (2017) View-Independent Working Memory Representations of Artificial Shapes in Prefrontal and Posterior Regions of the Human Brain. Cereb Cortex:1–16.

Christophel TB, Cichy RM, Hebart MN, Haynes J-D (2015) Parietal and early visual cortices encode working memory content across mental transformations. NeuroImage 106:198–206.

Curtis CE, D'Esposito M (2003) Persistent activity in the prefrontal cortex during working memory. Trends Cogn Sci 7:415–423.

Ester EF, Sprague TC, Serences JT (2015) Parietal and Frontal Cortex Encode Stimulus-Specific Mnemonic Representations during Visual Working Memory. Neuron 87:893–905.

Foster JJ, Sutterer DW, Serences JT, Vogel EK, Awh E (2016) The topography of alpha-band activity tracks the content of spatial working memory. J Neurophysiol 115:168–177.

García-Pérez MA (1998) Forced-choice staircases with fixed step sizes: asymptotic and small-sample properties. Vision Res 38:1861–1881.

Gayet S, Guggenmos M, Christophel TB, Haynes J-D, Paffen CLE, Stigchel SV der, Sterzer P (2017) Visual working memory enhances the neural response to matching visual input. J Neurosci:3418–16.

Grootswagers T, Wardle SG, Carlson TA (2016) Decoding Dynamic Brain Patterns from Evoked Responses: A Tutorial on Multivariate Pattern Analysis Applied to Time Series Neuroimaging Data. J Cogn Neurosci 29:677–697.

Harrison SA, Tong F (2009) Decoding reveals the contents of visual working memory in early visual areas. Nature 458:632–635.

Haufe S, Meinecke F, Görgen K, Dähne S, Haynes J-D, Blankertz B, Bießmann F (2014) On the interpretation of weight vectors of linear models in multivariate neuroimaging. NeuroImage 87:96–110.

Haxby JV, Connolly AC, Guntupalli JS (2014) Decoding Neural Representational Spaces Using Multivariate Pattern Analysis. Annu Rev Neurosci 37:435–456.

Hebb DO (1968) Concerning Imagery. In: Images, Perception, and Knowledge, pp 139–153 The University of Western Ontario Series in Philosophy of Science. Springer, Dordrecht. Available at: https://link.springer.com/chapter/10.1007/978-94-010-1193-8_7 [Accessed July 19, 2017].

Jerde TA, Merriam EP, Riggall AC, Hedges JH, Curtis CE (2012) Prioritized Maps of Space in Human Frontoparietal Cortex. J Neurosci 32:17382–17390.

King J-R, Dehaene S (2014) Characterizing the dynamics of mental representations: the temporal generalization method. Trends Cogn Sci 18:203–210.

King J-R, Pescetelli N, Dehaene S (2016) Brain Mechanisms Underlying the Brief Maintenance of Seen and Unseen Sensory Information. Neuron 92:1122–1134.

Kleiner M, Brainard D, Pelli D, Ingling A, Murray R, Broussard C, others (2007) What's new in Psychtoolbox-3. Perception 36:1.

Kok P, Mostert P, Lange FP de (2017) Prior expectations induce prestimulus sensory templates. Proc Natl Acad Sci:201705652.

Laeng B, Bloem IM, D'Ascenzo S, Tommasi L (2014) Scrutinizing visual images: The role of gaze in mental imagery and memory. Cognition 131:263–283.

Laeng B, Teodorescu D-S (2002) Eye scanpaths during visual imagery reenact those of perception of the same visual scene. Cogn Sci 26:207–231.

Manahova ME, Mostert P, Kok P, Schoffelen J-M, Lange FP de (2017) Stimulus familiarity and expectation jointly modulate neural activity in the visual ventral stream. bioRxiv:192518.

Maris E, Oostenveld R (2007) Nonparametric statistical testing of EEG- and MEG-data. J Neurosci Methods 164:177–190.

Mostert P, Kok P, de Lange FP (2015) Dissociating sensory from decision processes in human perceptual decision making. Sci Rep 5:18253.

Oostenveld R, Fries P, Maris E, Schoffelen J-M (2010) FieldTrip: Open Source Software for Advanced Analysis of MEG, EEG, and Invasive Electrophysiological Data. Comput Intell Neurosci 2011:e156869.

Plöchl M, Ossandón JP, König P (2012) Combining EEG and eye tracking: identification, characterization, and correction of eye movement artifacts in electroencephalographic data. Front Hum Neurosci 6 Available at: http://www.ncbi.nlm.nih.gov/pmc/articles/PMC3466435/.

Pratte MS, Tong F (2014) Spatial specificity of working memory representations in the early visual cortex. J Vis 14:22–22.

Rademaker RL, Serences JT (2017) Pinging the brain to reveal hidden memories. Nat Neurosci 20:767–769.

Rose NS, LaRocque JJ, Riggall AC, Gosseries O, Starrett MJ, Meyering EE, Postle BR (2016) Reactivation of latent working memories with transcranial magnetic stimulation. Science 354:1136–1139.

Serences JT, Ester EF, Vogel EK, Awh E (2009) Stimulus-Specific Delay Activity in Human Primary Visual Cortex. Psychol Sci 20:207–214.

Spaak E, Watanabe K, Funahashi S, Stokes MG (2017) Stable and Dynamic Coding for Working Memory in Primate Prefrontal Cortex. J Neurosci 37:6503–6516.

Spivey MJ, Geng JJ (2001) Oculomotor mechanisms activated by imagery and memory: Eye movements to absent objects. Psychol Res 65:235–241.

Sreenivasan KK, Curtis CE, D'Esposito M (2014) Revisiting the role of persistent neural activity during working memory. Trends Cogn Sci 18:82–89.

Stokes MG (2015) “Activity-silent” working memory in prefrontal cortex: a dynamic coding framework. Trends Cogn Sci 19:394–405.

Stokes MG, Kusunoki M, Sigala N, Nili H, Gaffan D, Duncan J (2013) Dynamic Coding for Cognitive Control in Prefrontal Cortex. Neuron 78:364–375.

Van Veen BD, van Drongelen W, Yuchtman M, Suzuki A (1997) Localization of brain electrical activity via linearly constrained minimum variance spatial filtering. IEEE Trans Biomed Eng 44:867–880.

Wolff MJ, Ding J, Myers NE, Stokes MG (2015) Revealing hidden states in visual working memory using electroencephalography. Front Syst Neurosci:123.

Wolff MJ, Jochim J, Akyürek EG, Stokes MG (2017) Dynamic hidden states underlying working-memory-guided behavior. Nat Neurosci advance online publication Available at: http://www.nature.com/neuro/journal/vaop/ncurrent/full/nn.4546.html [Accessed May 10, 2017].

